# Differential expression of striatal proteins in a mouse model of DOPA-responsive dystonia reveals shared mechanisms among dystonic disorders

**DOI:** 10.1101/2021.01.12.426367

**Authors:** Maria A. Briscione, Ashok R. Dinasarapu, Pritha Bagchi, Yuping Donsante, Anthony M. Downs, Xueliang Fan, Jessica Hoehner, H.A. Jinnah, Ellen J. Hess

## Abstract

Dystonia is characterized by involuntary muscle contractions that cause debilitating twisting movements and postures. Although basal ganglia dysfunction is implicated in many forms of dystonia, the underlying mechanisms are unclear. Therefore, to reveal abnormal striatal cellular processes and pathways implicated in dystonia, we used an unbiased proteomic approach in a knockin mouse model of DOPA-responsive dystonia, a model in which the striatum is known to play a central role in the expression of dystonia. Fifty-seven of the 1805 proteins identified were differentially regulated in DOPA-responsive dystonia mice compared to control mice. Most differentially regulated proteins were associated with gene ontology terms that implicated either mitochondrial or synaptic dysfunction whereby proteins associated with mitochondrial function were generally over-represented whereas proteins associated with synaptic function were largely under-represented. Remarkably, nearly 20% of the differentially regulated proteins identified in our screen are associated with pathogenic variants that cause inherited dystonic disorders in humans suggesting shared mechanisms across many different forms of dystonia.

## Introduction

Dystonia is characterized by involuntary muscle contractions that cause debilitating twisting movements and postures [1]. Dystonia is a heterogeneous disorder that may be sporadic or inherited [2, 3] and can sometimes occur as a result of brain injury, such as the dystonia that frequently occurs in cerebral palsy patients. Although the etiologies are diverse, basal ganglia dysfunction is consistently implicated across many different forms of dystonia [4]. Lesions and structural defects of the basal ganglia and its connections are sometimes accompanied by dystonia [5-8]. Functional imaging studies have revealed abnormal metabolic activity in the basal ganglia [9-12] and dystonia improves in some patients after neurosurgical lesions or deep brain stimulation of the internal segment of the globus pallidus [13, 14]. Additionally, dopamine (DA) neurotransmission, which is integral to the normal function of the basal ganglia, is abnormal in both sporadic and inherited forms of dystonia. Mutations in genes critical for the synthesis of DA, including GTP-cyclohydrolase and tyrosine hydroxylase (*TH*) cause DOPA-responsive dystonia [DRD; 15, 16, 17]. Idiopathic forms of dystonia such as writer’s cramp and spasmodic dysphonia are also associated with abnormal striatal dopaminergic neurotransmission [18, 19]. Further, reduced striatal D2 dopamine receptor (D2R) availability is observed in both inherited and idiopathic forms of dystonia including DYT1 dystonia, blepharospasm, torticollis, writer’s cramp and laryngeal dystonia [18-23]. Despite the overwhelming evidence implicating basal ganglia dysfunction in dystonia, the precise nature of the cellular defects that give rise to dystonia are unclear.

To provide insight into the striatal cellular processes and pathways underlying dystonia, we used an unbiased proteomic approach in a mouse knockin model of DRD. DRD is caused by mutations in genes necessary for the synthesis of DA, which mediates striatal activity [24, 25], and is therefore a prototype for understanding basal ganglia dysfunction in dystonia. DRD knockin mice carry the human DRD-causing p.381Q>K mutation in *TH* and exhibit the characteristic features of DRD including reduced brain DA concentrations and dystonic movements that improve in response to L-DOPA administration [26]. Further, it is known that striatal DA neurotransmission plays a central role in mediating the dystonia in DRD mice [26]. Our unbiased assessment of striatal protein expression in DRD mice compared to control mice revealed 57 proteins that were differentially expressed. Remarkably, pathogenic variants of 12 of the 57 differentially regulated proteins are associated with disorders in which dystonia is a prominent feature, suggesting shared mechanisms across many different forms of dystonia.

## Methods

### Mice

Male and female adult (3-5 month) mice homozygous for the c.1160C>A *TH* mutation (*Th^drd^*/*Th^drd^* ;DRD mice) and normal littermates (+/+) were used for all experiments. The DRD mutation is coisogenic on C57BL/6J and congenic on DBA/2J. DRD and normal mice were generated as F1 hybrids of C57BL/6J +/*Th^drd^* x DBA/2J +/*Th^drd^* to circumvent the high perinatal lethality exhibited by inbred C57BL/6J DRD mice. Cell-type specific reporter transgenes inbred on C57BL/6J were bred onto the C57BL/6J +/*Th^drd^* strain prior to the F1 hybrid cross to identify D1R-expressing (*Drd1a*-tdTomato; JAX) or D2R-expressing (*Drd2*-EGFP; Heintz, Rockefeller University) spiny projection neurons (SPNs); [27, 28]. Mice were maintained as described by Rose et al.[26], housed in standard mouse cages (2-5 mice/cage) in standard environmental conditions (72°C, 40-50% relative humidity) with a 12 hr light cycle (7am to 7 pm) and *ad lib* access to food and water. All procedures conformed to the NIH Guidelines for the Care and Use of Animals and were approved by the Emory University Animal Care and Use Committee.

### Behavioral assessment

Mice were habituated to the test cages (29 x 50 cm) for > 3 hrs prior to behavioral assessment. A behavioral inventory was used to identify abnormal movements including tonic flexion (forelimbs, hindlimbs, trunk, head), tonic extension (forelimbs, hindlimbs, trunk, head), clonus (forelimbs and hindlimbs), twisting (trunk, head), and tremor [forelimbs, hindlimbs, trunk, head; 29]. Testing began at 2 pm, and abnormal movements were observed and scored by an observer blinded to genotype for 30 sec at 10 min intervals for 1 hr. Abnormal movements were scored as present (1) or absent (0) for each body region during each time bin. An abnormal movement score was calculated by summing all scores.

### Immunohistochemistry

Mice were deeply anesthetized with isoflurane and perfused with 4% ice-cold, buffered paraformaldehyde (pH 7.2), incubated overnight in the paraformaldehyde perfusate at 4°C and transferred to a 30% buffered sucrose solution. Brains were sectioned (30 μm) in the coronal plane using a freezing microtome. Sections were immunostained using anti-mCherry primary antibody (1:5,000; #AB167453; Abcam) or GFP polyclonal primary antibody (1:20,000; A-11122; Invitrogen). Floating sections were treated with 0.5% Triton-X in Tris-buffered saline (TBS) for 30 min and then transferred to 5% normal goat serum, 5% bovine serum albumin, 0.1% bovine gelatin, 0.05% Tween-80, and 0.01% sodium azide in TBS for 2 hrs. Sections were incubated with primary antibody for 16-24 hrs at 4°C and then for 2 hrs at 4°C with biotinylated goat anti-rabbit IgG secondary antibody (1:800, Vector Laboratories) in 5% normal goat serum in 0.1% Triton-X-100 in TBS. Sections were incubated with avidin-biotin complex (Vector Labs, Burlingame, CA) for 1 hr, and developed using 3-3’-diaminobenzidine (DAB; Sigma-Aldrich, St. Louis, MO) as the chromogen.

### Image analysis

QuPath, an open source software for whole slide image analysis, was used to automate counts of immunoreactive cells and to define striatal regions for analysis. Striatal sections from Bregma +1.145 to 0.145, based on the Allen Coronal Mouse Brain Atlas, were quantified. Striatal regions were defined by outlining and calculating the entire area of the striatum. To consistently define dorsomedial, dorsolateral, and ventral striatum across sections and mice, the center x and y coordinate of the right and left striatum was determined using QuPath. A line between the striatal midpoints delimited the dorsal and ventral striatum and a line perpendicular to the dorsal-ventral boundary at the center x,y coordinate defined medial and lateral striatum. To determine cell counts for the dorsomedial (DM), dorsolateral (DL), and ventral striatum, the total number of positive cells from each region was divided by total area of the region to determine the number of positive cells/mm^2^.

### Preparation of tissue for proteomic analysis

Mice were euthanized by cervical dislocation. Brains were rapidly removed and striata were rapidly dissected and frozen on dry ice and stored at −80° C. The protocol for tissue homogenization was adapted from a published method [30]. Samples (∼30 mg) were vortexed in 300 µL of urea lysis buffer (8 M urea, 10 mM Tris, 100 mM NaH2PO4, pH 8.5), including 3 µL (100x stock) HALT(-EDTA) protease and phosphatase inhibitor cocktail (Pierce). All homogenization was performed using a Bullet Blender (Next Advance) according to the manufacturer’s protocol. Briefly, each tissue piece was added to urea lysis buffer in a 1.5 mL Rino tube (Next Advance) harboring 750 mg stainless steel beads (0.9-2 mm in diameter) and blended twice for 5 minute intervals in the cold room (4° C). Protein homogenates were transferred to 1.5 mL Eppendorf tubes and were sonicated (Sonic Dismembrator, Fisher Scientific) 3 times for 5 sec each with 15 sec intervals of rest at 30% amplitude to disrupt nucleic acids and were subsequently centrifuged at 4° C . Protein concentration was determined by the bicinchoninic acid (BCA) method, and samples were frozen in aliquots at −80 °C. Protein homogenates (100 µg) were diluted with 50 mM NH4HCO3 to a final concentration of less than 2 M urea and then were treated with 1 mM dithiothreitol (DTT) at room temperature for 30 min, followed by 5 mM iodoacetimide at room temperature for 30 min in the dark. Protein samples were digested with 1:100 (w/w) lysyl endopeptidase (Wako) at room temperature for 4 hrs and were further digested overnight with 1:50 (w/w) trypsin (Promega) at room temperature. Resulting peptides were desalted with HLB column (Waters) and were dried under vacuum.

### Proteomics data acquisition

The data acquisition by LC-MS/MS protocol was adapted from a published procedure[30] and was provided and completed by the Integrated Proteomics Core, an Emory Neuroscience NINDS Core Facilities (ENNCF): www.cores.emory.edu/eipc/resources/index.html. Derived peptides were resuspended in loading buffer (0.1% trifluoroacetic acid). Peptide mixtures (3 µL) were separated on a self-packed C18 (1.9 µm Dr. Maisch, Germany) fused silica column (25 cm x 75 µM internal diameter (ID); New Objective, Woburn, MA) by a Dionex Ultimate 3000 RSLCNano and monitored on a Fusion mass spectrometer (ThermoFisher Scientific, San Jose, CA). Elution was performed over a 140 min gradient at a rate of 300 nL/min with buffer B ranging from 3% to 99% (buffer A: 0.1% formic acid in water, buffer B: 0.1 % formic acid in acetonitrile). The mass spectrometer cycle was programmed to collect at the top speed for 3 sec cycles. The MS scans (300-1500 m/z range, 200,000 AGC target, 50 ms maximum injection time) were collected at a resolution of 120,000 at m/z 200 in profile mode and the HCD MS/MS spectra (1.5 m/z isolation width, 30% collision energy, 10,000 AGC target, 35 ms maximum injection time) were detected in the ion trap. Dynamic exclusion was set to exclude previous sequenced precursor ions for 20 sec within a 10 ppm window. Precursor ions with +1, and +8 or higher charge states were excluded from sequencing.

### Protein identification and quantification

ThermoFisher generated RAW files were used for label-free quantitation (LFQ) of proteins in eight biological samples. Base peak chromatograms were inspected visually using RawMeat software. All RAW files were processed together in a single run by MaxQuant version 1.6.0.16 with default parameters unless otherwise specified (http://www.maxquant.org). Database searches were performed using the Andromeda search engine (a peptide search engine based on probabilistic scoring) with the UniProt mouse sequence database (update in 2017). MaxQuant provides a contaminants.fasta database file (a database of common laboratory contaminants) within the software that is automatically added to the list of proteins for the in-silico digestion when this feature is enabled.

Precursor mass tolerance was set to 4.5 ppm in the main search, and fragment mass tolerance was set to 20 ppm. Digestion enzyme specificity was set to trypsin with a maximum of 2 missed cleavages. A minimum peptide length of 6 residues was required for identification. Up to 5 modifications per peptide were allowed; acetylation (protein N-terminal), oxidation (Met) and deamidation (NQ) were set as variable modifications, and carbamidomethylation (Cys) was set as a fixed modification. No Andromeda score threshold was set for unmodified peptides. A minimum Andromeda score of 40 was required for modified peptides. Peptide and protein false discovery rates (FDR) were both set to 1% based off a target-decoy reverse database. Proteins that shared all identified peptides were combined into a single protein group. If all identified peptides from one protein were a subset of identified peptides from another protein, these proteins were also combined into that group. Peptides that matched multiple protein groups (“razor” peptides) were assigned to the protein group with the most unique peptides.

Peaks were detected in Full MS, and a three-dimensional peak was constructed as a function of peak centroid m/z (7.5 ppm threshold) and peak area over time. Following de-isotoping, peptide intensities were determined by extracted ion chromatograms based on the peak area at the retention time with the maximum peak height. Peptide intensities were normalized to minimize overall proteome difference based on the assumption that most peptides do not change in intensity between samples. Protein LFQ intensities were calculated from the median of pairwise intensity ratios of peptides identified in two or more samples and adjusted to the cumulative intensity across samples. Quantification was performed using razor and unique peptides, including those modified by acetylation (protein N-terminal), oxidation (Met) and deamidation (NQ). A minimum peptide ratio of 1 was required for protein intensity normalization, and “Fast LFQ” was enabled.

### Data analysis

Data analysis was performed using Perseus version 1.5.0.31 (http://www.perseus-framework.org). Contaminants and protein groups identified by a single peptide were filtered from the data set. FDR was calculated as the percentage of reverse database matches out of total forward and reverse matches. Protein group LFQ intensities were log2 transformed to reduce the effect of outliers. Data were filtered by valid values. A minimum of four valid values was required in at least one group (DRD or normal). For cluster analysis and statistical comparisons between proteomes, protein groups missing LFQ values were assigned values using imputation. Missing values were assumed to be biased toward low abundance proteins that were below the MS detection limit. The missing values were replaced with random values taken from a median downshifted Gaussian distribution to simulate low abundance LFQ values. Imputation was performed separately for each sample from a distribution with a width of 0.3 and downshift of 1.8. Hierarchical clustering was performed on Z-score normalized, log2 LFQ intensities using Euclidean distance and average linkage with k-means preprocessing (300 clusters). Log-fold changes were calculated as the difference in log2 LFQ intensity averages between experimental and control groups. Welsch’s t test calculations were used in statistical tests as histograms of LFQ intensities showed that all data sets approximated normal distributions. Protein IDs/Protein names were mapped to gene symbols before further steps, Heatmap and volcano plots were created using R Statistical Software (Foundation for Statistical Computing, Vienna, Austria).

### Gene Ontology (GO) and KEGG pathway enrichment analysis of the differentially expressed proteins

Analyses were performed using the gene list of differentially expressed proteins (p <0.05) and annotations were limited by species (m*us musculus).* DAVID Bioinformatic Resources v6.8 (https://david.ncifcrf.gov) [31] was used to identify KEGG pathways (Kyoto Encyclopedia of Genes and Genomes). The Gene Ontology Resource (http://geneontology.org) Panther Overrepresentation Test (released 20200407; GO database released 20200323) was used to identify enriched biological terms using the Fisher Exact test and FDR correction [32, 33].

### Identification of dystonia-associated proteins

PubMed searches were performed to determine whether proteins differentially regulated in DRD mice compared to normal mice (p < 0.05) were previously associated with dystonia. Keywords included the differentially regulated protein (protein and gene name) and “dystonia”. Studies were reviewed and only those that were clearly associated with both dystonia and the protein of interest were included. When differentially regulated proteins were components of a larger protein complex, the larger protein complex was also used as a keyword.

## Results

### Striatal anatomy

The development of the striatum is, in part, mediated by DA. Therefore, we first determined if anatomical features of the striatum in DRD mice differed from normal dopamine-intact mice because potential anatomical abnormalities caused by the loss of DA in developing DRD mice would influence the interpretation of proteomic analyses. There was no apparent difference in the size of the striatum between normal and DRD mice. The vast majority of neurons in the striatum are spiny projections neurons (SPNs). Generally, there are two subtypes of SPNs, D1 dopamine receptor (D1R)-expressing SPNs and D2R-expressing SPNs, which each express a unique repertoire of proteins. To determine if the SPN subtype populations in DRD mice differed from normal mice, we assessed the number and distribution of D1R-expressing or D2R-expression SPNs by using reporter transgenes that express tdTomato (tdTom) in D1R-SPNs or EGFP in D2R-SPNs to identify each subtype. The density of tdTom- and EGFP-labeled cells within the striatum was not significantly different between normal and DRD mice (Figure 1). Further, the distribution of EGFP- and tdTom-positive neurons in dorsomedial, dorsolateral, and ventral striatum in DRD mice did not differ from normal mice (Figure 1). To ensure that the reporter transgenes did not spuriously alter the dystonic phenotype of DRD mice, which is known to be dependent on striatal dysfunction, we assessed abnormal movements in DRD mice carrying tdTom (n=6), EGFP (n=5) or both tdTom and EGFP BAC transgenes (n=7). Abnormal movements in DRD mice carrying the transgenes were not significantly different from DRD mice without the transgenes (not shown; n=27; Student’s t test, p>0.1 for all). These results suggest that major anatomical features of the striatum are intact in DRD mice.

**Figure 1.**
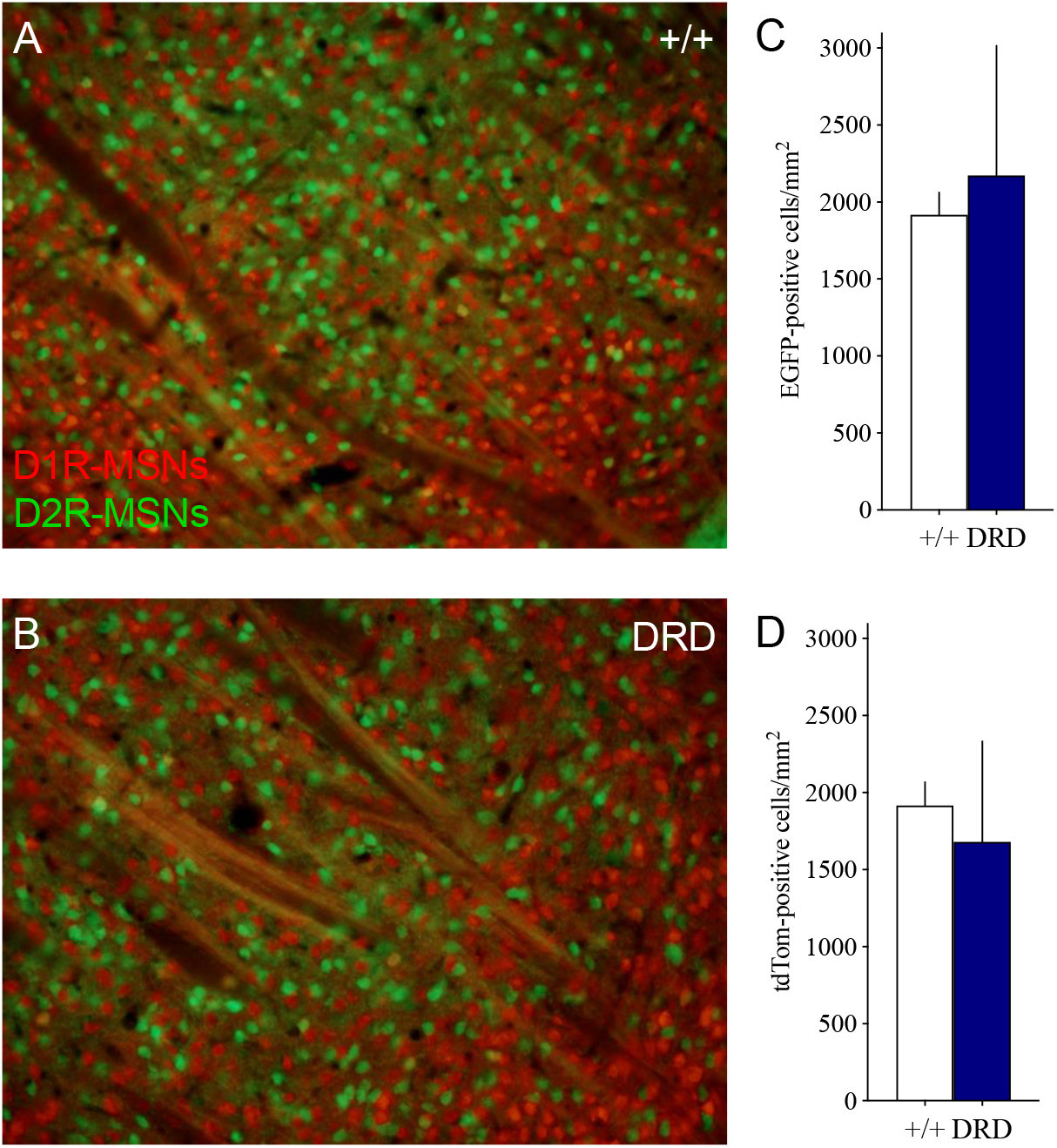
Representative fluorescent micrographs of DRD and normal mouse striatum. Representative micrographs of coronal sections from normal (**A**) and DRD (**B**) mouse striatum with endogenous tdTom (D1R-MSNs) and EGFP (D2R-MSNs) fluorescent proteins. **C**. EGFP positive cell counts (D2R-MSNs) did not differ between DRD (n=3) and normal (n=4) mice (p>0.1; Student’s *t* test). **D**. tdTom positive cell counts (D1R-MSNs) did not differ between DRD (n=3) and normal (n=3) mice (p>0.1; Student’s *t* test). Values represent mean + SEM.

### Proteomic analyses

The unbiased examination of the striatal proteome identified 1805 proteins. These proteins are illustrated in a volcano plot (Figure 2) which shows the magnitude of differences on the X-axis and the unadjusted p-value on the Y-axis. Proteins that are reduced in DRD mice compared to control mice fall to the left of zero, and those increased in DRD mice are to the right of zero. TH protein expression was significantly reduced in DRD mice compared to normal mice (Figure 2) and reached a false discovery rate of p<0.05. This is consistent with our previous work that demonstrates a reduction in TH protein content and activity that is caused by the knockin point mutation in DRD mice [26].

**Figure 2.**
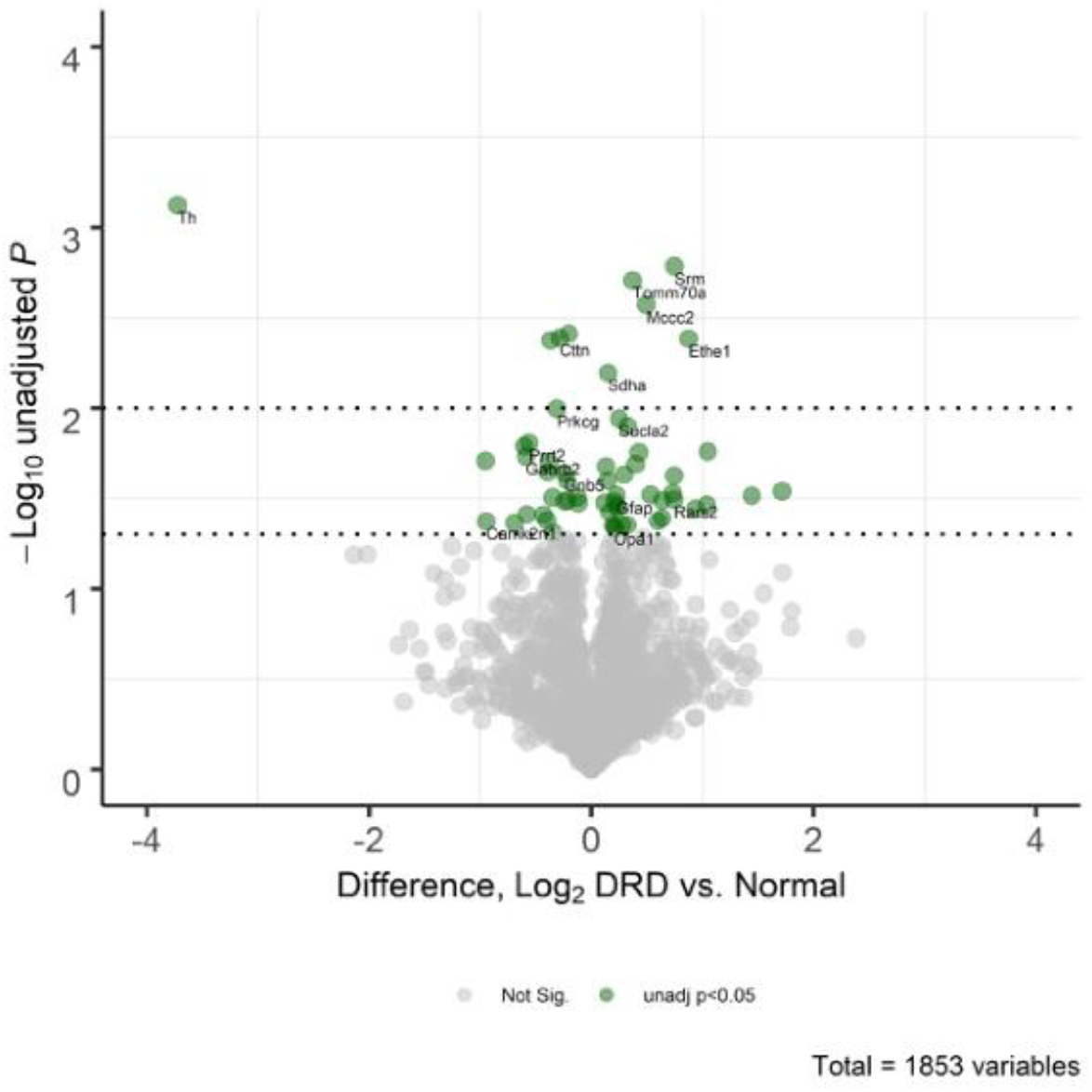
Volcano plot of differentially expressed proteins in normal and DRD mice. Each dot corresponds to an identified protein, including those that do not have associated gene names. The y-axis shows negative log10 transformed p-values obtained based on a two-tailed Welch’s t-tests with dotted lines representing unadjusted p-values of 0.05 and <0.01. The x-axis illustrates the log2 difference between DRD and normal mouse striatal protein expression with negative values representing proteins that are reduced in DRD mice compared to normal mice. Green dots indicate proteins with unadjusted p-values < 0.05; those previously associated with dystonia are labeled with the associated gene name.

The number of differentially expressed protein identified depends on statistical thresholds and methods. Further, it has been suggested that the false discovery rate (FDR) is too conservative for exploratory proteomics[34]. Therefore, results are presented using several different approaches including uncorrected p-values and volcano plots. We identified proteins differentially regulated in DRD compared to normal mice using the following arbitrary p-values cutoffs: p<0.001, 0.01 and 0.05. TH was significantly downregulated in DRD mice at p<0.001. Eight additional proteins were differentially regulated in the striatum of DRD compared to normal mice at p<0.01. When p<0.05 was considered, 19 additional proteins were downregulated and 29 additional proteins were upregulated in the striatum of DRD compared to normal mice (Figure 3a). Differentially regulated proteins (p<0.05) are illustrated in a heatmap (Figure 3b).

**Figure 3.**
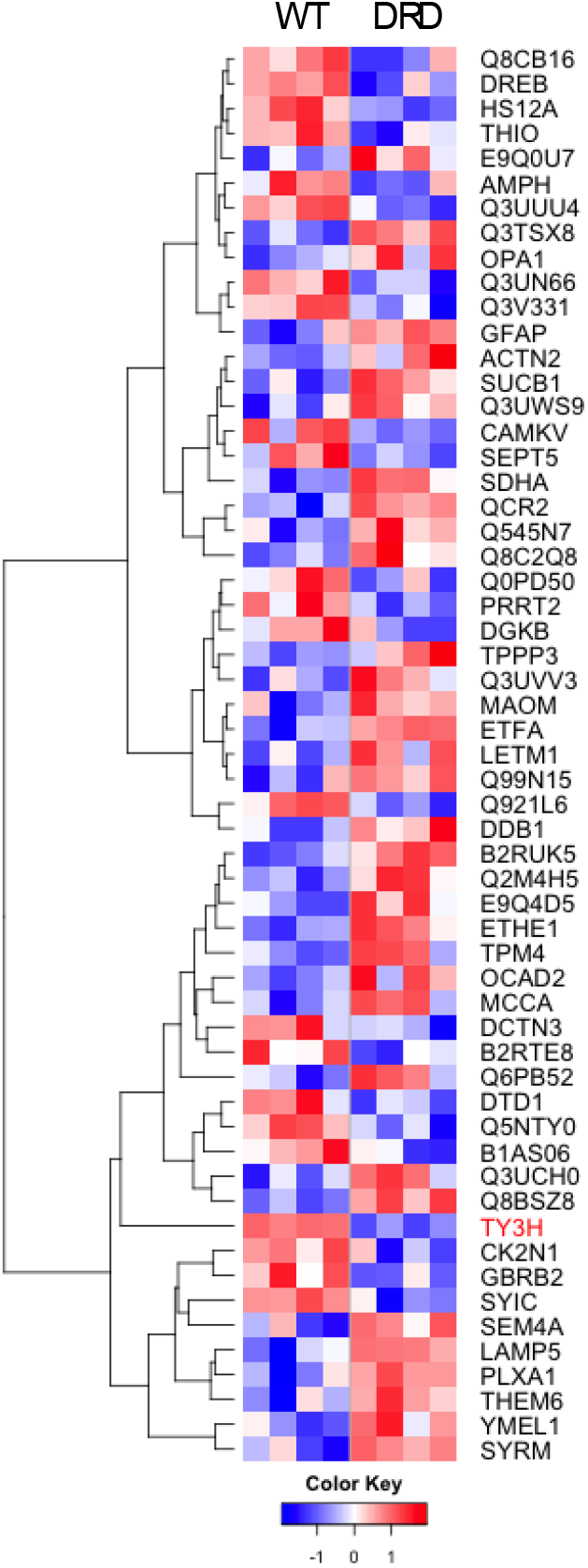
Heat map of differentially expressed proteins in the striatum of normal and DRD mice. Relative levels of differentially regulated proteins (p<0.05) are presented. Rows represent proteins and columns represent samples. Proteins clustered based on protein expression (abundance) pattern. Low to high protein expression is represented by a change of color from blue to red, respectively. The color key scale bar at bottom shows z-score values for the heatmap. Each row (protein) is scaled to have mean zero and standard deviation one.

Gene ontology analysis was used to identify pathways and functions that may be abnormally regulated in DRD mice. When differentially regulated proteins with p values < 0.05 were considered, several dysregulated functional groups were identified (Gene Ontology FDR p<0.005 for all). Table 1 provides a representative sample of the gene ontologies with nonspecific terms such as ‘cytoplasm’ or ‘cell projection’ or terms that included < 4 differentially expressed proteins omitted for clarity. Two prominent categories of gene ontologies were identified: mitochondrion and synapse. Proteins associated with the gene ontology term ‘mitochondrion’ accounted for ∼36% of the differentially expressed proteins. Mitochondrial subcategories included both structural and functional (reactome) terms including mitochondrial membrane or matrix and mitochondrial energetics (TCA cycle). All differentially-expressed proteins associated with these gene ontology terms were over-represented in DRD mice compared to normal mice. Proteins associated with the gene ontology term ‘synapse’ accounted for ∼33% of the differentially expressed proteins. In contrast to the mitochondrial-associated proteins, differentially expressed proteins associated with synaptic function, including pre- and postsynapse, synaptic vesicle, axon and glutamatergic synapse, were under-represented in DRD mice compared to normal mice with the exception of Lamp5, Rpl6 and Actn2.

**Table 1.** Gene Ontology (GO) enrichment analysis of differentially regulated proteins

| GO term | Genes | p value | FDR |
| --- | --- | --- | --- |
| Mitochondrion | Sucla2\|Sdha\|Rars2\|Mccc1\|Prdx5\|Txn\|Hsd17b10\|Ociad2\|Dnaja1\|Atp5c1\|Opa1\|Ethe1\|Yme1l1\|Me2\|Th\|Uqcrc2\|Letm1\|Ckmt1\|Etfa\|Tomm70a\|Mccc2 | 1.11E-09 | 1.10E-06 |
| Synapse | Camk2n1\|Prkcg\|Lamp5\|Amph\|Dlgap3\|Palm\|Cttn\|Dbn1\|Rpl6\|Gabrb2\|Sept5\|Camkv\|Gnb5\|Th\|Actn2\|Prrt2\|Dgkb\|Rab8a\|Sept6 | 5.74E-09 | 3.80E-06 |
| Myelin Sheath | Omg\|Sucla2\|Sdha\|Atp5c1\|Gfap\|Gnb5\|Uqcrc2\|Ckmt1 | 1.52E-07 | 7.56E-05 |
| Postsynapse | Camk2n1\|Prkcg\|Dlgap3\|Palm\|Cttn\|Dbn1\|Rpl6\|Gabrb2\|Camkv\|Actn2\|Prrt2\|Rab8a | 6.59E-07 | 1.63E-04 |
| Mitochondrial Membrane | Sdha\|Mccc1\|Hsd17b10\|Ociad2\|Atp5c1\|Opa1\|Yme1l1\|Uqcrc2\|Letm1\|Ckmt1\|Tomm70a | 1.02E-06 | 2.03E-04 |
| Postsynaptic Specialization | Camk2n1\|Prkcg\|Dlgap3\|Palm\|Dbn1\|Rpl6\|Rab8a\|Gabrb2\|Actn2 | 3.17E-06 | 4.50E-04 |
| Axon | Prkcg\|Lamp5\|Txn\|Amph\|Palm\|Cttn\|Dbn1\|Sept5\|Th\|Prrt2\|Sept6 | 3.89E-06 | 5.15E-04 |
| Glutamatergic Synapse | Amph\|Dlgap3\|Cttn\|Dbn1\|Camkv\|Actn2\|Prrt2\|Dgkb\|Rab8a | 1.06E-05 | 1.05E-03 |
| Synaptic Vesicle | Amph\|Sept5\|Th\|Prrt2\|Sept6\|Rab8a | 9.60E-05 | 7.06E-03 |
| ATP Binding | Sucla2\|Rars2\|Mccc1\|Prkcg\|Srm\|Dnaja1\|Yme1l1\|Camkv\|Hsph1\|Ckmt1\|Iars\|Hspa12a\|Dgkb\|Mccc2 | 1.98E-05 | 7.65E-03 |
| tRNA Binding | Hsd17b10\|Dtd1\|Iars\|Rpl6 | 3.20E-05 | 1.06E-02 |
| Presynapse | Prkcg\|Amph\|Sept5\|Gnb5\|Th\|Prrt2\|Rab8a\|Sept6 | 2.18E-04 | 1.24E-02 |
| Ligase Activity | Sucla2\|Mccc1\|Iars\|Rars2\|Mccc2 | 5.81E-05 | 1.41E-02 |
| The citric acid cycle (TCA) and respiratory electron transport | Sucla2\|Sdha\|Atp5c1\|Opa1\|Me2\|Etfa | 3.78E-05 | 2.08E-02 |
| Dendrite | Camk2n1\|Prkcg\|Txn\|Dlgap3\|Opa1\|Dbn1\|Th\|Rab8a | 6.94E-04 | 3.36E-02 |
| Cellular response to stress | Prdx5\|Txn\|Dnaja1\|Dctn3\|Hsph1\|Hspa12a | 1.40E-04 | 3.85E-02 |
| Mitochondrial Matrix | Mccc1\|Hsd17b10\|Uqcrc2\|Mccc2\|Etfa | 9.72E-04 | 4.02E-02 |

DAVID was used to probe for novel signaling pathways within the KEGG pathway database that may be differentially regulated in DRD compared to normal mouse striatum. Eight enriched KEGG pathways were identified (Table 2), including GABAergic and endocannabinoid signaling pathways. In addition, the Parkinson’s disease pathway was identified. Both Parkinson’s disease and DRD are characterized by dopamine deficiency, a central biochemical feature of both disorders, providing additional support of the validity of this proteomic analysis.

**Table 2.** Enriched KEGG pathways of differentially expressed proteins

| Pathway Description | Genes | p value | FDR q-value |
| --- | --- | --- | --- |
| Parkinson's disease | Uqcrc2\|Sept5\|Sdha\|Th\|Atp5c1 | 0.003 | 0.259453884 |
| Valine, leucine, and isoleucine degradation | Mccc2\|Hsd17b10\|Mccc1 | 0.022 | 0.643639237 |
| Metabolic pathways | Uqcrc2\|Sdha\|Th\|Atp5c1\|Mccc2\|Hsd17b10\|Mccc1\|Dgkb\|SRM\|Ckmt1\|Sucla2 | 0.029 | 0.598691922 |
| Alzheimer's disease | UQCRC2\|SDHA\|ATP5C1\|HSD17B10 | 0.036 | 0.582326059 |
| GABAergic synapse | Gabrb2\|Gnb5\|Prkcg | 0.050 | 0.621737675 |
| Morphine addiction | Gabrb2\|Gnb5\|Prkcg | 0.057 | 0.599647403 |
| Retrograde endocannabinoid signaling | Gabrb2\|Gnb5\|Prkcg | 0.068 | 0.611250528 |
| Carbon metabolism | Me2\|Sdha\|Sucla2 | 0.083 | 0.640892686 |

A literature review of the 57 differentially regulated proteins (p<0.05) revealed 12 proteins in which pathogenic genetic variants are associated with dystonia (Table 3). Thus, ∼20% of the differentially regulated proteins in DRD mice are implicated in other forms of dystonia.

**Table 3.**
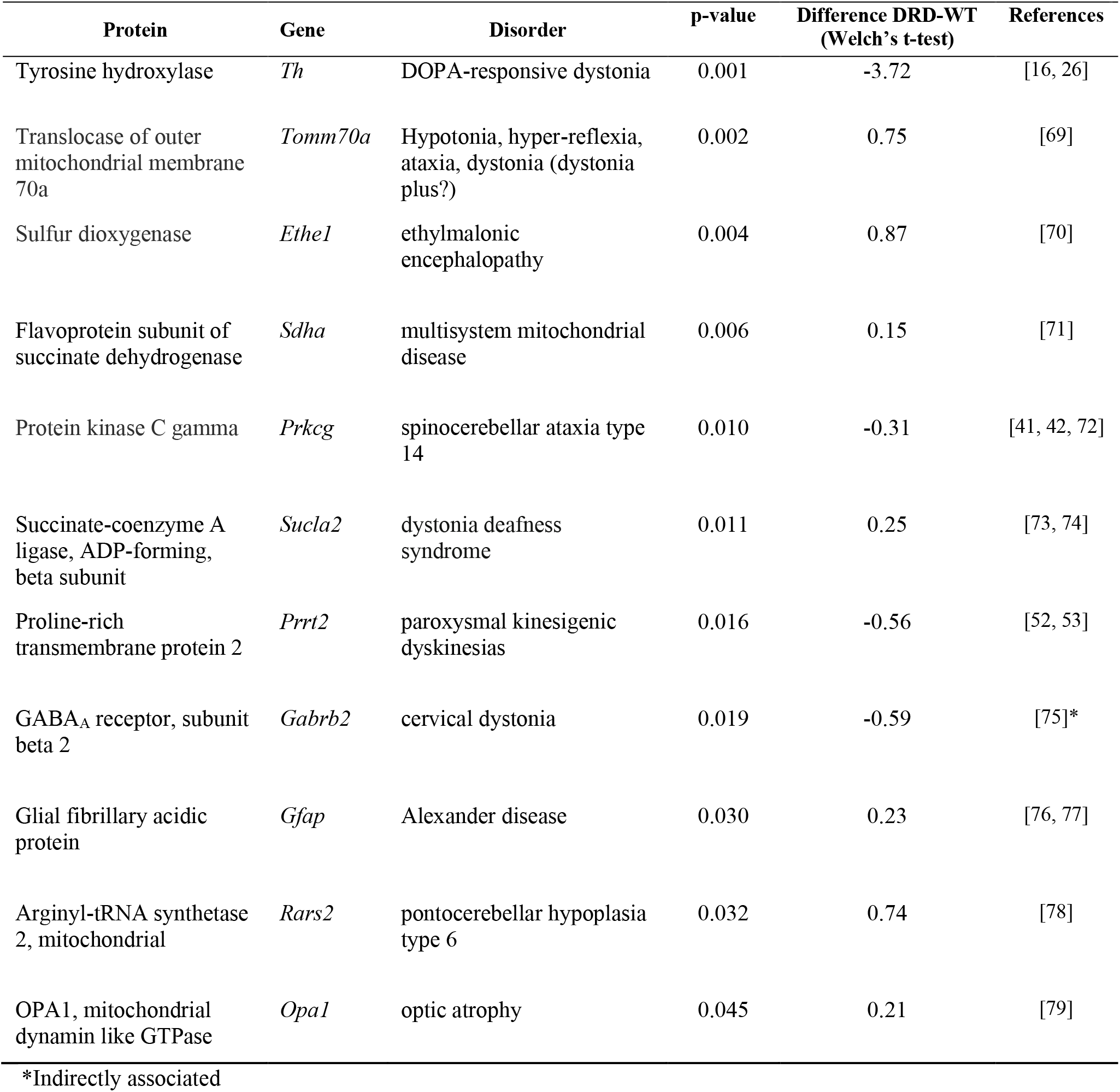
Differentially regulated proteins previously associated with dystonia

## Discussion

Dystonia is generally not associated with degeneration or overt cellular pathology, suggesting that neuronal dysfunction mediates the abnormal movements. Therefore, to better understand the cellular defects underlying dystonia, we examined striatal protein expression in a mouse model of DRD because it is known that the dystonic movements in these mice are mediated by striatal dysfunction.[26] Our unbiased assessment of the DRD mouse striatal proteome revealed that of the 1805 proteins identified in the proteomic screen only 57 (∼3%) were differentially regulated in DRD mice compared to normal mice. Importantly, the proteomic analysis revealed that TH protein expression was significantly reduced in DRD mice. This finding is consistent with our previous anatomical and biochemical analyses that demonstrated a reduction in TH expression and activity [26], providing validation for the results of the proteomic analysis.

Despite the significant deficit in TH expression that is caused by the mutation in *Th*, major anatomical features of the striatum, including striatal size and the distribution and number of D1R- and D2R-expressing SPNs, were intact in DRD mice. These results suggest that the proteomic analysis was not overtly confounded by gross morphological defects. That relatively few proteins were abnormally regulated is consistent with the observation that striatal anatomy is preserved in DRD mice. However, it is surprising that we did not observe more extensive protein dysregulation considering that, in adults, dopamine neurotransmission plays an important role in the strength and plasticity of glutamatergic synaptic inputs, the intrinsic excitability of SPNs and contributes to morphological changes[24, 35, 36]. Dopamine neurotransmission also plays a critical role in striatal development. DA innervates the nascent striatum mid-gestation, prior to the arrival of excitatory afferents and induces the formation of immature spines through both D1Rs and D2Rs[37, 38]. DA also regulates the macroscopic dendritic architecture of SPNs by stimulating neurite outgrowth in embryonic striatal neurons through activation of D1Rs but not D2Rs[39, 40], suggesting that an early-life DA deficit may reduce dendritic complexity. Subtle morphological changes such as spine formation and complexity would not have been detected in our gross anatomical analyses but were nonetheless reflected in the proteomic analysis.

Many of the differentially regulated proteins were associated with synaptic function, which is perhaps not surprising considering that the brain is enriched in these proteins. It is, however, striking that the majority (16/19) of the differentially expressed proteins represented in the GO ‘synapse’ pathway and related pathways were significantly under-represented in DRD mice, suggesting a general deficit in striatal neurotransmission. Specifically, the intracellular signaling protein kinase C gamma (Prkcg) and the transmitter release protein proline-rich-transmembrane-protein-2 (Prrt2) are notable because of their shared dysregulation across dystonias. Prkcg is a neuron-specific calcium-activated, phospholipid- and diacylglycerol (DAG)-dependent serine/threonine-protein kinase. Although variants in Prkcg are specifically associated with the dystonic disorder spinocerebellar ataxia type 14 [41-43], identification of Prkcg in our proteomic analysis has broader implications for dystonia in light of the role of Prkcg in the regulation of synaptic plasticity. Abnormal plasticity is observed in many different forms of dystonia in humans and in several mouse models of dystonia [44-50], suggesting that Prkcg may be a pathophysiological node. Prrt2 is a presynaptic membrane protein that interacts with SNAP-25 to mediate calcium-dependent vesicular exocytosis, particularly glutamate release [51]. Pathogenic variants in *PRRT2* are associated with paroxysmal kinesigenic dyskinesia, a disorder that includes dystonic movements [52-54]. Prrt2 dysregulation is also observed in a mouse model of Dyt6 (*Thap1*). Because both Prkcg and Prrt2 are associated with specific forms of dystonia and are also more broadly implicated in dystonias, these proteins are attractive targets for therapeutics aimed at enhancing expression and/or activity.

Approximately one-third of all differentially regulated proteins were associated with synaptic function based on the enriched GO terms. Subsets of the broad term ‘synapse’ implicated some specific functions, like ‘presynapse,’ which was expected considering that DRD is caused by a mutation in TH, which is located primarily in nigrostriatal terminals. Consistent with a presynaptic dopamine defect in DRD, the KEGG analysis identified Parkinson’s disease, which is, in part, caused by striatal dopamine deficiency that results from degeneration of nigrostriatal neurons. Enriched pathways for glutamatergic (GO) and GABAergic (KEGG) synapses were also identified. The striatum receives abundant glutamatergic innervation from both corticostriatal and thalamostriatal efferents while the vast majority of neurons in the striatum itself are GABAergic spiny projection neurons. The activity of both is mediated by dopamine neurotransmission. Thus, the enriched pathways reflect both the biochemical defect and striatal anatomy, supporting the validity of the pathway analyses.

Approximately one-third of all differentially regulated proteins were associated with mitochondrial function based on the enriched GO terms. Further, fully half of the proteins associated with dystonia in Table 3 play a role in mitochondrial function including *Ethe1, Sdha, Sucla2, Rars2, Opa1*, and *Tomm70a*. Several additional lines of evidence implicate mitochondrial dysfunction across different forms of dystonia. In addition to the inherited dystonias listed in Table 3, other inherited dystonias in which mitochondrial dysfunction is implicated include [55, 56] Leber’s hereditary optic neuropathy which is caused by a mutation in the mtDNA complex I gene *MTND6* [57] and deafness-dystonia-optic neuropathy (*DDON*) syndrome which is caused by a mutation in the deafness-dystonia peptide (*DDP1*) gene that encodes the mitochondrial translocase subunit Tim8 A [58]. Mitochondrial dysfunction is also implicated in idiopathic dystonias. In particular, a reduction in mitochondrial complex I activity was observed in patients with idiopathic focal [59, 60], segmental and generalized dystonia [59]. Dystonia can also be induced by accidental exposure to the mitochondrial complex II inhibitor 3-nitropropionic acid (3-NPA) in otherwise healthy individuals [61-63]. Thus, mitochondrial dysfunction is a shared defect among inherited, acquired, and idiopathic dystonias in humans. Furthermore, mitochondrial dysregulation is also shared among mouse models, including Dyt1 (*Tor1a*), Dyt6 (*Thap1*), and now DRD (*Th*), demonstrating that mitochondrial changes can occur even when the dystonia-causing defect is apparently unrelated to mitochondrial function [64, 65]. However, it will be critical to determine whether the mitochondrial abnormalities identified here are integral to the expression of dystonia or compensatory.

Pathogenic variants of twelve of the 57 differentially regulated proteins identified in DRD mice are implicated in other forms of dystonia. That approximately 20% of the differentially regulated proteins are also associated with rare inherited dystonias suggests common pathological mechanisms across different forms of dystonia. Abundant evidence implicating abnormal dopaminergic neurotransmission in many different dystonias has been accumulating for decades. More recently, in addition to the shared mechanisms such as mitochondrial and synaptic dysfunction mentioned previously, abnormal plasticity, developmental defects and cholinergic dysfunction have been implicated across different forms of dystonias using -omics, computational or physiological approaches in both humans and mouse models[64-68]. The identification of shared processes suggests that it may be possible to identify specific, druggable targets for the development of novel therapeutics that are effective in a broad range of dystonias.

## Acknowledgements

We thank Alec P. Shannon for technical support. This work was supported by the National Institutes of Health, National Institute of Neurological Disorders and Stroke Grant R01 NS088528.

## Notes

The authors report no financial conflicts of interest concerning the research reported in this manuscript.

### Competing Interest Statement

The authors have declared no competing interest.

## References

1. Albanese A, Bhatia K, Bressman SB, Delong MR, Fahn S, Fung VS, Hallett M, Jankovic J, Jinnah HA, Klein C, Lang AE, Mink JW and Teller JK (2013) Phenomenology and classification of dystonia: a consensus update. Mov Disord 28:863–73. doi: 10.1002/mds.25475

2. Jinnah HA and Sun YV (2019) Dystonia genes and their biological pathways. Neurobiol Dis 129:159–168. doi: 10.1016/j.nbd.2019.05.014

3. Balint B, Mencacci NE, Valente EM, Pisani A, Rothwell J, Jankovic J, Vidailhet M and Bhatia KP (2018) Dystonia. Nat Rev Dis Primers 4:25. doi: 10.1038/s41572-018-0023-6

4. Neychev VK, Gross RE, Lehericy S, Hess EJ and Jinnah HA (2011) The functional neuroanatomy of dystonia. Neurobiol Dis 42:185–201. doi: S0969-9961(11)00047-7 [pii] 10.1016/j.nbd.2011.01.026

5. Marsden CD, Obeso JA, Zarranz JJ and Lang AE (1985) The anatomical basis of symptomatic hemidystonia. Brain 108:463–83.

6. Pettigrew LC and Jankovic J (1985) Hemidystonia: a report of 22 patients and a review of the literature. J Neurol Neurosurg Psychiatry 48:650–7.

7. Obeso JA and Gimenez-Roldan S (1988) Clinicopathological correlation in symptomatic dystonia. Adv Neurol 50:113–22.

8. Bhatia KP and Marsden CD (1994) The behavioural and motor consequences of focal lesions of the basal ganglia in man. Brain 117:859–76.

9. Kerrison JB, Lancaster JL, Zamarripa FE, Richardson LA, Morrison JC, Holck DE, Andreason KW, Blaydon SM and Fox PT (2003) Positron emission tomography scanning in essential blepharospasm. Am J Ophthalmol 136:846–52. doi: S000293940300895X [pii]

10. Galardi G, Perani D, Grassi F, Bressi S, Amadio S, Antoni M, Comi GC, Canal N and Fazio F (1996) Basal ganglia and thalamo-cortical hypermetabolism in patients with spasmodic torticollis. Acta Neurol Scand 94:172–176.

11. Carbon M, Niethammer M, Peng S, Raymond D, Dhawan V, Chaly T, Ma Y, Bressman S and Eidelberg D (2009) Abnormal striatal and thalamic dopamine neurotransmission: Genotype-related features of dystonia. Neurology 72:2097–103. doi: 10.1212/WNL.0b013e3181aa538f

12. Eidelberg D, Moeller J R., Antonini A, Kazumata K, Nakamura T, Dhawan V, Spetsieris P, deLeon D, Bressman S, B. and Fahn S (1998) Functional brain networks in DYT1 dystonia. Ann Neurol 44:303–312.

13. Centen LM, Oterdoom DLM, Tijssen MAJ, Lesman-Leegte I, van Egmond ME and van Dijk JMC (2020) Bilateral Pallidotomy for Dystonia: A Systematic Review. Mov Disord. doi: 10.1002/mds.28384

14. Rodrigues FB, Duarte GS, Prescott D, Ferreira J and Costa J (2019) Deep brain stimulation for dystonia. Cochrane Database Syst Rev 1:CD012405. doi: 10.1002/14651858.CD012405.pub2

15. Ichinose H, Ohye T, Takahashi E, Seki N, Hori T, Segawa M, Nomura Y, Endo K, Tanaka H, Tsuji S, Fujita K and Nagatsu T (1994) Hereditary progressive dystonia with marked diurnal fluctuation caused by mutations in the GTP cyclohydrolase I gene. Nat Genet 8:236–242.

16. Knappskog PM, Flatmark T, Mallet J, Ludecke B and Bartholome K (1995) Recessively inherited L-DOPA-responsive dystonia caused by a point mutation (Q381K) in the tyrosine hydroxylase gene. Human Molecular Genetics 4:1209–1212.

17. van den Heuvel LP, Luiten B, Smeitink JA, de Rijk-van Andel JF, Hyland K, Steenbergen-Spanjers GC, Janssen RJ and Wevers RA (1998) A common point mutation in the tyrosine hydroxylase gene in autosomal recessive L-DOPA-responsive dystonia in the Dutch population. Hum Genet 102:644–6.

18. Berman BD, Hallett M, Herscovitch P and Simonyan K (2013) Striatal dopaminergic dysfunction at rest and during task performance in writer’s cramp. Brain 136:3645–58. doi: 10.1093/brain/awt282

19. Simonyan K, Berman BD, Herscovitch P and Hallett M (2013) Abnormal striatal dopaminergic neurotransmission during rest and task production in spasmodic dysphonia. J Neurosci 33:14705–14. doi: 10.1523/JNEUROSCI.0407-13.2013

20. Horie C, Suzuki Y, Kiyosawa M, Mochizuki M, Wakakura M, Oda K, Ishiwata K and Ishii K (2009) Decreased dopamine D2 receptor binding in essential blepharospasm. Acta Neurol Scand 119:49–54. doi: ANE1053 [pii] 10.1111/j.1600-0404.2008.01053.x

21. Hierholzer J, Cordes M, Schelosky L, Richter W, Keske U, Venz S, Semmler W, Poewe W and Felix R (1994) Dopamine D2 receptor imaging with iodine-123-iodobenzamide SPECT in idiopathic rotational torticollis. J Nucl Med 35:1921–7.

22. Naumann M, Pirker W, Reiners K, Lange KW, Becker G and Brucke T (1998) Imaging the pre- and postsynaptic side of striatal dopaminergic synapses in idiopathic cervical dystonia: a SPECT study using [123I] epidepride and [123I] beta-CIT. Mov Disord 13:319–23. doi: 10.1002/mds.870130219

23. Asanuma K, Ma Y, Okulski J, Dhawan V, Chaly T, Carbon M, Bressman SB and Eidelberg D (2005) Decreased striatal D2 receptor binding in non-manifesting carriers of the DYT1 dystonia mutation. Neurology 64:347–9. doi: 64/2/347 [pii] 10.1212/01.WNL.0000149764.34953.BF

24. Gerfen CR and Surmeier DJ (2011) Modulation of striatal projection systems by dopamine. Annu Rev Neurosci 34:441–66. doi: 10.1146/annurev-neuro-061010-113641

25. DeLong MR and Wichmann T (2007) Circuits and circuit disorders of the basal ganglia. Arch Neurol 64:20–4. doi: 10.1001/archneur.64.1.20

26. Rose SJ, Yu XY, Heinzer AK, Harrast P, Fan XL, Raike RS, Thompson VB, Pare JF, Weinshenker D, Smith Y, Jinnah HA and Hess EJ (2015) A new knock-in mouse model of L-DOPA-responsive dystonia. Brain 138:2987–3002. doi: 10.1093/brain/awv212

27. Ade KK, Wan Y, Chen M, Gloss B and Calakos N (2011) An Improved BAC Transgenic Fluorescent Reporter Line for Sensitive and Specific Identification of Striatonigral Medium Spiny Neurons. Front Syst Neurosci 5:32. doi: 10.3389/fnsys.2011.00032

28. Chan CS, Peterson JD, Gertler TS, Glajch KE, Quintana RE, Cui Q, Sebel LE, Plotkin JL, Shen W, Heiman M, Heintz N, Greengard P and Surmeier DJ (2012) Strain-specific regulation of striatal phenotype in Drd2-eGFP BAC transgenic mice. J Neurosci 32:9124–32. doi: 10.1523/JNEUROSCI.0229-12.2012

29. Raike RS, Pizoli CE, Weisz C, van den Maagdenberg AM, Jinnah HA and Hess EJ (2013) Limited regional cerebellar dysfunction induces focal dystonia in mice. Neurobiol Dis 49:200–10. doi: 10.1016/j.nbd.2012.07.019

30. Seyfried NT, Dammer EB, Swarup V, Nandakumar D, Duong DM, Yin L, Deng Q, Nguyen T, Hales CM, Wingo T, Glass J, Gearing M, Thambisetty M, Troncoso JC, Geschwind DH, Lah JJ and Levey AI (2017) A Multi-network Approach Identifies Protein-Specific Co-expression in Asymptomatic and Symptomatic Alzheimer’s Disease. Cell Syst 4:60–72 e4. doi: 10.1016/j.cels.2016.11.006

31. Huang da W, Sherman BT and Lempicki RA (2009) Systematic and integrative analysis of large gene lists using DAVID bioinformatics resources. Nat Protoc 4:44–57. doi: 10.1038/nprot.2008.211

32. Ashburner M, Ball CA, Blake JA, Botstein D, Butler H, Cherry JM, Davis AP, Dolinski K, Dwight SS, Eppig JT, Harris MA, Hill DP, Issel-Tarver L, Kasarskis A, Lewis S, Matese JC, Richardson JE, Ringwald M, Rubin GM and Sherlock G (2000) Gene ontology: tool for the unification of biology. The Gene Ontology Consortium. Nat Genet 25:25–9. doi: 10.1038/75556

33. The Gene Ontology C (2019) The Gene Ontology Resource: 20 years and still GOing strong. Nucleic Acids Res 47:D330–D338. doi: 10.1093/nar/gky1055

34. Pascovici D, Handler DC, Wu JX and Haynes PA (2016) Multiple testing corrections in quantitative proteomics: A useful but blunt tool. Proteomics 16:2448–53. doi: 10.1002/pmic.201600044

35. Day M, Wang Z, Ding J, An X, Ingham CA, Shering AF, Wokosin D, Ilijic E, Sun Z, Sampson AR, Mugnaini E, Deutch AY, Sesack SR, Arbuthnott GW and Surmeier DJ (2006) Selective elimination of glutamatergic synapses on striatopallidal neurons in Parkinson disease models. Nat Neurosci 9:251–9. doi: 10.1038/nn1632

36. Villalba RM and Smith Y (2013) Differential striatal spine pathology in Parkinson’s disease and cocaine addiction: A key role of dopamine? Neuroscience 251:2–20. doi: 10.1016/j.neuroscience.2013.07.011

37. Fasano C, Bourque MJ, Lapointe G, Leo D, Thibault D, Haber M, Kortleven C, Desgroseillers L, Murai KK and Trudeau LE (2013) Dopamine facilitates dendritic spine formation by cultured striatal medium spiny neurons through both D1 and D2 dopamine receptors. Neuropharmacology 67:432–43. doi: 10.1016/j.neuropharm.2012.11.030

38. Money KM and Stanwood GD (2013) Developmental origins of brain disorders: roles for dopamine. Front Cell Neurosci 7:260. doi: 10.3389/fncel.2013.00260

39. Schmidt U, Pilgrim C and Beyer C (1998) Differentiative effects of dopamine on striatal neurons involve stimulation of the cAMP/PKA pathway. Mol Cell Neurosci 11:9–18. doi: 10.1006/mcne.1998.0668

40. Schmidt U, Beyer C, Oestreicher AB, Reisert I, Schilling K and Pilgrim C (1996) Activation of dopaminergic D1 receptors promotes morphogenesis of developing striatal neurons. Neuroscience 74:453–60.

41. Ganos C, Zittel S, Minnerop M, Schunke O, Heinbokel C, Gerloff C, Zuhlke C, Bauer P, Klockgether T, Munchau A and Baumer T (2014) Clinical and neurophysiological profile of four German families with spinocerebellar ataxia type 14. Cerebellum 13:89–96. doi: 10.1007/s12311-013-0522-7

42. Miura S, Nakagawara H, Kaida H, Sugita M, Noda K, Motomura K, Ohyagi Y, Ayabe M, Aizawa H, Ishibashi M and Taniwaki T (2009) Expansion of the phenotypic spectrum of SCA14 caused by the Gly128Asp mutation in PRKCG. Clin Neurol Neurosurg 111:211–5. doi: 10.1016/j.clineuro.2008.09.013

43. Nibbeling EA, Delnooz CC, de Koning TJ, Sinke RJ, Jinnah HA, Tijssen MA and Verbeek DS (2017) Using the shared genetics of dystonia and ataxia to unravel their pathogenesis. Neurosci Biobehav Rev 75:22–39. doi: 10.1016/j.neubiorev.2017.01.033

44. Quartarone A, Bagnato S, Rizzo V, Siebner HR, Dattola V, Scalfari A, Morgante F, Battaglia F, Romano M and Girlanda P (2003) Abnormal associative plasticity of the human motor cortex in writer’s cramp. Brain 126:2586–96. doi: 10.1093/brain/awg273

45. Edwards MJ, Huang YZ, Mir P, Rothwell JC and Bhatia KP (2006) Abnormalities in motor cortical plasticity differentiate manifesting and nonmanifesting DYT1 carriers. Mov Disord 21:2181–6. doi: 10.1002/mds.21160

46. Evinger C (2015) Benign Essential Blepharospasm is a Disorder of Neuroplasticity: Lessons From Animal Models. J Neuroophthalmol 35:374–9. doi: 10.1097/WNO.0000000000000317

47. Gilbertson T, Humphries M and Steele JD (2019) Maladaptive striatal plasticity and abnormal reward-learning in cervical dystonia. Eur J Neurosci. doi: 10.1111/ejn.14414

48. Martella G, Tassone A, Sciamanna G, Platania P, Cuomo D, Viscomi MT, Bonsi P, Cacci E, Biagioni S, Usiello A, Bernardi G, Sharma N, Standaert DG and Pisani A (2009) Impairment of bidirectional synaptic plasticity in the striatum of a mouse model of DYT1 dystonia: role of endogenous acetylcholine. Brain 132:2336–49. doi: awp194 [pii] 10.1093/brain/awp194

49. Weise D, Schramm A, Beck M, Reiners K and Classen J (2011) Loss of topographic specificity of LTD-like plasticity is a trait marker in focal dystonia. Neurobiol Dis 42:171–6. doi: 10.1016/j.nbd.2010.11.009

50. Meunier S, Russmann H, Shamim E, Lamy JC and Hallett M (2012) Plasticity of cortical inhibition in dystonia is impaired after motor learning and paired-associative stimulation. Eur J Neurosci 35:975–86. doi: 10.1111/j.1460-9568.2012.08034.x

51. Li M, Niu F, Zhu X, Wu X, Shen N, Peng X and Liu Y (2015) PRRT2 Mutant Leads to Dysfunction of Glutamate Signaling. Int J Mol Sci 16:9134–51. doi: 10.3390/ijms16059134

52. Marano M, Motolese F, Consoli F, De Luca A and Di Lazzaro V (2018) Paroxysmal Dyskinesias in a PRRT2 Mutation Carrier. Tremor Other Hyperkinet Mov (N Y) 8:616. doi: 10.7916/D8S488X0

53. Zhang Y, Li L, Chen W, Gan J and Liu ZG (2017) Clinical characteristics and PRRT2 gene mutation analysis of sporadic patients with paroxysmal kinesigenic dyskinesia in China. Clin Neurol Neurosurg 159:25–28. doi: 10.1016/j.clineuro.2017.05.004

54. Bruno MK, Ravina B, Garraux G, Hallett M, Ptacek L, Singleton A, Johnson J, Hanson M, Considine E and Gwinn-Hardy K (2004) Exercise-induced dystonia as a preceding symptom of familial Parkinson’s disease. Mov Disord 19:228–30. doi: 10.1002/mds.10626

55. De Vries DD, Went LN, Bruyn GW, Scholte HR, Hofstra RM, Bolhuis PA and van Oost BA (1996) Genetic and biochemical impairment of mitochondrial complex I activity in a family with Leber hereditary optic neuropathy and hereditary spastic dystonia. Am J Hum Genet 58:703–11.

56. Sarzi E, Brown MD, Lebon S, Chretien D, Munnich A, Rotig A and Procaccio V (2007) A novel recurrent mitochondrial DNA mutation in ND3 gene is associated with isolated complex I deficiency causing Leigh syndrome and dystonia. Am J Med Genet A 143A:33–41. doi: 10.1002/ajmg.a.31565

57. Huoponen K, Vilkki J, Aula P, Nikoskelainen EK and Savontaus ML (1991) A new mtDNA mutation associated with Leber hereditary optic neuroretinopathy. Am J Hum Genet 48:1147–53.

58. Roesch K, Curran SP, Tranebjaerg L and Koehler CM (2002) Human deafness dystonia syndrome is caused by a defect in assembly of the DDP1/TIMM8a-TIMM13 complex. Hum Mol Genet 11:477–86. doi: 10.1093/hmg/11.5.477

59. Benecke R, Strumper P and Weiss H (1992) Electron transfer complex I defect in idiopathic dystonia. Ann Neurol 32:683–6. doi: 10.1002/ana.410320512

60. Schapira AH, Warner T, Gash MT, Cleeter MW, Marinho CF and Cooper JM (1997) Complex I function in familial and sporadic dystonia. Ann Neurol 41:556–9. doi: 10.1002/ana.410410421

61. Ming L (1995) Moldy sugarcane poisoning--a case report with a brief review. J Toxicol Clin Toxicol 33:363.7.

62. He F, Zhang S, Qian F and Zhang C (1995) Delayed dystonia with striatal CT lucencies induced by a mycotoxin (3-nitropropionic acid). Neurology 45:2178–83.

63. Peraica M, Radic B, Lucic A and Pavlovic M (1999) Toxic effects of mycotoxins in humans. Bull World Health Organ 77:754–66.

64. Zakirova Z, Fanutza T, Bonet J, Readhead B, Zhang W, Yi Z, Beauvais G, Zwaka TP, Ozelius LJ, Blitzer RD, Gonzalez-Alegre P and Ehrlich ME (2018) Mutations in THAP1/DYT6 reveal that diverse dystonia genes disrupt similar neuronal pathways and functions. PLoS Genet 14:e1007169. doi: 10.1371/journal.pgen.1007169

65. Martin JN, Bair TB, Bode N, Dauer WT and Gonzalez-Alegre P (2009) Transcriptional and proteomic profiling in a cellular model of DYT1 dystonia. Neuroscience 164:563–72. doi: 10.1016/j.neuroscience.2009.07.068

66. Mencacci NE, Reynolds R, Ruiz SG, Vandrovcova J, Forabosco P, Sanchez-Ferrer A, Volpato V, Consortium UKBE, International Parkinson’s Disease Genomics C, Weale ME, Bhatia KP, Webber C, Hardy J, Botia JA and Ryten M (2020) Dystonia genes functionally converge in specific neurons and share neurobiology with psychiatric disorders. Brain 143:2771–2787. doi: 10.1093/brain/awaa217

67. Frederick NM, Shah PV, Didonna A, Langley MR, Kanthasamy AG and Opal P (2019) Loss of the dystonia gene Thap1 leads to transcriptional deficits that converge on common pathogenic pathways in dystonic syndromes. Hum Mol Genet 28:1343–1356. doi: 10.1093/hmg/ddy433

68. Eskow Jaunarajs KL, Scarduzio M, Ehrlich ME, McMahon LL and Standaert DG (2019) Diverse Mechanisms Lead to Common Dysfunction of Striatal Cholinergic Interneurons in Distinct Genetic Mouse Models of Dystonia. J Neurosci 39:7195–7205. doi: 10.1523/JNEUROSCI.0407-19.2019

69. Dutta D, Briere LC, Kanca O, Marcogliese PC, Walker MA, High FA, Vanderver A, Krier J, Carmichael N, Callahan C, Taft RJ, Simons C, Helman G, Network UD, Wangler MF, Yamamoto S, Sweetser DA and Bellen HJ (2020) De novo mutations in TOMM70, a receptor of the mitochondrial import translocase, cause neurological impairment. Hum Mol Genet 29:1568–1579. doi: 10.1093/hmg/ddaa081

70. Walsh DJ, Sills ES, Lambert DM, Gregersen N, Malone FD and Walsh AP (2010) Novel ETHE1 mutation in a carrier couple having prior offspring affected with ethylmalonic encephalopathy: Genetic analysis, clinical management and reproductive outcome. Mol Med Rep 3:223–6. doi: 10.3892/mmr_00000243

71. Renkema GH, Wortmann SB, Smeets RJ, Venselaar H, Antoine M, Visser G, Ben-Omran T, van den Heuvel LP, Timmers HJ, Smeitink JA and Rodenburg RJ (2015) SDHA mutations causing a multisystem mitochondrial disease: novel mutations and genetic overlap with hereditary tumors. Eur J Hum Genet 23:202–9. doi: 10.1038/ejhg.2014.80

72. Nibbeling EAR, Duarri A, Verschuuren-Bemelmans CC, Fokkens MR, Karjalainen JM, Smeets C, de Boer-Bergsma JJ, van der Vries G, Dooijes D, Bampi GB, van Diemen C, Brunt E, Ippel E, Kremer B, Vlak M, Adir N, Wijmenga C, van de Warrenburg BPC, Franke L, Sinke RJ and Verbeek DS (2017) Exome sequencing and network analysis identifies shared mechanisms underlying spinocerebellar ataxia. Brain 140:2860–2878. doi: 10.1093/brain/awx251

73. Garone C, Gurgel-Giannetti J, Sanna-Cherchi S, Krishna S, Naini A, Quinzii CM and Hirano M (2017) A Novel SUCLA2 Mutation Presenting as a Complex Childhood Movement Disorder. J Child Neurol 32:246–250. doi: 10.1177/0883073816666221

74. Maas RR, Marina AD, de Brouwer AP, Wevers RA, Rodenburg RJ and Wortmann SB (2016) SUCLA2 Deficiency: A Deafness-Dystonia Syndrome with Distinctive Metabolic Findings (Report of a New Patient and Review of the Literature). JIMD Rep 27:27–32. doi: 10.1007/8904_2015_464

75. Berman BD, Pollard RT, Shelton E, Karki R, Smith-Jones PM and Miao Y (2018) GABAA Receptor Availability Changes Underlie Symptoms in Isolated Cervical Dystonia. Front Neurol 9:188. doi: 10.3389/fneur.2018.00188

76. Jefferson RJ, Absoud M, Jain R, Livingston JH, MS Vdk and Jayawant S (2010) Alexander disease with periventricular calcification: a novel mutation of the GFAP gene. Dev Med Child Neurol 52:1160–3. doi: 10.1111/j.1469-8749.2010.03784.x

77. Machol K, Jankovic J, Vijayakumar D, Burrage LC, Jain M, Lewis RA, Fuller GN, Xu M, Penas-Prado M, Gule-Monroe MK, Rosenfeld JA, Chen R, Eng CM, Yang Y, Lee BH, Moretti PM, Undiagnosed Diseases N and Dhar SU (2018) Atypical Alexander disease with dystonia, retinopathy, and a brain mass mimicking astrocytoma. Neurol Genet 4:e248. doi: 10.1212/NXG.0000000000000248

78. Glamuzina E, Brown R, Hogarth K, Saunders D, Russell-Eggitt I, Pitt M, de Sousa C, Rahman S, Brown G and Grunewald S (2012) Further delineation of pontocerebellar hypoplasia type 6 due to mutations in the gene encoding mitochondrial arginyl-tRNA synthetase, RARS2. J Inherit Metab Dis 35:459–67. doi: 10.1007/s10545-011-9413-6

79. Liskova P, Ulmanova O, Tesina P, Melsova H, Diblik P, Hansikova H, Tesarova M and Votruba M (2013) Novel OPA1 missense mutation in a family with optic atrophy and severe widespread neurological disorder. Acta Ophthalmol 91:e225–31. doi: 10.1111/aos.12038

